# QUANTIFYING ACYL CHAIN INTERDIGITATION IN SIMULATED BILAYERS VIA DIRECT TRANSBILAYER INTERACTIONS

**DOI:** 10.1101/2024.12.20.629658

**Authors:** Emily H. Chaisson, Frederick A. Heberle, Milka Doktorova

## Abstract

In a lipid bilayer, the interactions between lipid hydrocarbon chains from opposing leaflets can influence membrane properties. These interactions include the phenomenon of interdigitation, in which an acyl chain of one leaflet extends past the bilayer midplane and into the opposing leaflet. While static interdigitation is well understood in gel phase bilayers from X-ray diffraction measurements, much less is known about dynamic interdigitation in fluid phases. In this regard, atomistic molecular dynamics simulations can provide mechanistic information on interleaflet interactions that can be used to generate experimentally testable hypotheses. To address limitations of existing computational methodologies which provide results that are either indirect or averaged over time and space, here we introduce three novel ways of quantifying the extent of chain interdigitation. Our protocols include the analysis of instantaneous interactions at the level of individual carbon atoms, thus providing temporal and spatial resolution for a more nuanced picture of dynamic interdigitation. We compare the methods on bilayers composed of lipids with equal total number of carbon atoms but different mismatches between the sn-1 and sn-2 chain lengths. We find that these metrics, which are based on freely available software packages and are easy to implement, provide complementary details that help characterize various features of lipid-lipid contacts at the bilayer midplane. The new frameworks thus allow for a deeper look at fundamental molecular mechanisms underlying bilayer structure and dynamics, and present a valuable expansion of the membrane biophysics toolkit.

## INTRODUCTION

Biological membranes are chemically complex, consisting of a lipid bilayer with hundreds of chemically distinct lipids and various types of proteins that interact with them. The structure and physical behavior of the bilayer are determined in part by the structure and thermodynamic properties of its constituent lipids. As one example, lipids with two chains of varying lengths (i.e. asymmetric chain lipids), have been hypothesized to affect the fluidity of the membrane in both prokaryotic and eukaryotic cells.^1-7^ Asymmetry in the effective chain lengths of the lipids can arise both from unequal numbers of carbons as well as from the tilt of the glycerol backbone, which naturally allows the sn-1 chain to reach deeper into the hydrophobic core of the bilayer than the sn-2 chain. Thus, the resulting length mismatch within a lipid molecule can range from negligible (for some lipids with chains differing by 0 or 2 carbons^8^) to extreme, as in the case of some sphingomyelin species.^9^

A large mismatch in effective chain length increases the probability of interdigitation, a phenomenon where an acyl chain in one leaflet penetrates past the bilayer midplane into the opposing leaflet. In principle, enhanced interactions between the two leaflets in an interdigitated state could effectively ‘zip’ the leaflets together and thereby affect interleaflet friction. These interactions may also impact the overall flexibility of the bilayer and its mechanical properties although further studies and robust interdigitation metrics are needed to elucidate the exact relationship. Being an internal property of the lipid bilayer, interdigitation likely incurs an energy cost similar to those of lipid tilt and splay^10-14^ but the detailed form of the energy functional has yet to be determined.

Interdigitation in vitro has been studied with techniques such as X-ray and neutron diffraction^15-18^, differential scanning calorimetry^15, 17-19^, vibrational spectroscopy^15, 17^, nuclear magnetic resonance, electron microscopy and electron spin resonance^17^, scattering techniques^20, 21^, and fluorescence.^7, 17, 18, 22, 23^ Rather than directly quantifying interdigitation, these methods reveal its effects on membrane properties such as thickness, order, lateral lipid diffusion, and transition temperature. Though such indirect information is useful, many of these techniques suffer from poor spatial and/or temporal resolution, and may require extrinsic probes that can locally perturb the bilayer environment, potentially interfering with the measurement.^24^

Molecular dynamics (MD) simulations offer a complementary approach for characterizing membrane structure and dynamics by tracking the positions of individual lipid atoms over time. Interdigitation in simulated bilayers has previously been quantified in several ways: (1) from the overlap of mass density distributions for atoms (or atom groups) in opposing leaflets;^9, 18, 21, 25-27^ (2) by measuring the distance along the bilayer normal between terminal methyl carbons of lipids in opposing leaflets;^28, 29^ or (3) by measuring the changes in the position of individual atoms along the bilayer normal.^15^ For the latter, the underlying assumption is that transverse movement of these atoms brings the entire chain closer to or further away from the bilayer midplane and thus correlates with the extent of interdigitation. While these measurements constitute a reasonable starting point for interrogating interleaflet interactions, to our knowledge a detailed examination of pairwise contacts between lipid atoms from opposing leaflets has not yet been demonstrated.

Here, we develop three alternative ways of analyzing interdigitation in MD simulations to complement and expand on previous methods. Each new metric provides unique insight into different aspects of acyl chain interdigitation, such as specific trans-leaflet carbon-carbon interactions and the shape and evolution of the leaflet-leaflet interaction region. The three methods are: (1) **nISA** (<u>n</u>ormalized <u>I</u>nterfacial <u>S</u>urface <u>A</u>rea), which utilizes 3D Voronoi tessellations to define and quantify the interleaflet contact surface; (2) **ccMat** (<u>c</u>arbon <u>c</u>ontact <u>M</u>atrix), which fills a matrix with counts of the average number of direct interactions between carbon atom pairs in opposing leaflets; and (3) **doMat** (<u>d</u>ensity <u>o</u>verlap <u>M</u>atrix), which computes the overlap area of number density distributions for pairs of carbons in opposing leaflets. We benchmark these tools using two symmetric bilayers, 1,2-dipalmitoyl-sn-glycero-3-phosphocholine (16:0-16:0 PC, DPPC) and 1-myristoyl-2-stearoyl-sn-glycero-3-phosphocholine (14:0-18:0 PC, MSPC), that are expected to differ substantially in the extent of chain interdigitation.^21^

## METHODS

### Simulation details

When we started this work, 14:0-18:0 PC (MSPC) was not available in the CHARMM-GUI web server.^30^ To build this lipid, we first used CHARMM-GUI to construct a bilayer of 18:0-18:0-PC with 100 lipids per leaflet (200 lipids total) and hydrated with 45 waters per lipid without any salt ions. Following the protocol described in ref.^21^, 14:0-18:0 PC was then built by removing the terminal methyl and two adjoining methylene segments of the sn-1 chain (C316-C318 in CHARMM notation) as well as the two hydrogens on C315. A new terminal methyl, C314, was then created by converting C315 to hydrogen (H14Z) by changing its atom name, type, and partial charge. The two hydrogens on the C314 atom were also changed to match the atom name, type, and partial charge to reflect the characteristics of hydrogens in a terminal methyl group.

All simulations were run using the NAMD software^31^ and the CHARMM36 force field^32, 33^ for lipids. The MSPC bilayer was energy minimized for 1200 steps, then simulated for a total of 1 ns with an integration time-step of 1 fs before the production run, which used a 2 fs time-step. The DPPC bilayer was equilibrated using the CHARMM-GUI 6-step equilibration protocol. Simulations were run at a constant temperature of 50°C (323.15 K) and a constant pressure of 1 atm using NAMD’s Langevin thermostat and Nose-Hoover barostat, respectively. Long-range interactions were modeled with a 10−12 Å Lennard-Jones potential using NAMD’s *<u>vdwForceSwitching</u>* option. Hydrogen bonds were constrained with the *rigidbonds* parameter set to “all.” The Particle Mesh Ewald (PME) method was used and grid spacing was set to 1 Å to model the electrostatic interactions. An autocorrelation function from ref.^34^ was used to evaluate the equilibration of bilayer area; after convergence, each system was simulated for an additional ∼ 1 µs. The length of the analyzed trajectory was 0.991 µs for MSPC and 1.02 µs for DPPC. We also ran an additional NP*γ*T simulation starting from the end of the DPPC trajectory and applying a tension of *γ* = −5 mN/m in the *xy* plane which effectively compressed the bilayer area^35^ (Table S1). This simulation was run for a total of 0.4 µs with the last 0.36 µs used for analysis. Prior to analysis, all systems were centered in VMD^36^ by moving the center of mass of the lipid terminal methyl groups to *z* = 0. The top leaflet was then defined as the leaflet in which the headgroup phosphorus atom had an average *z* > 0.

### 3D Voronoi tessellation

Voronoi tessellations (VT) of simulated bilayers have previously been used to estimate areas and volumes of individual lipids.^37, 38^ Here, we used the atomic coordinates of lipid and water atoms as generators (i.e., the points used to construct Voronoi cells) to perform a 3D VT in each simulation frame using the software *voro++*.^39^ The Voronoi cells generated by the tessellation—each composed of a set of vertices, edges, and faces—represent the physical space occupied by individual atoms and enable various calculations as described below. We used a custom *Python* script to reformat the *voro++* output for subsequent analyses in MATLAB.^40^

### Normalized Interfacial Surface Area (nISA)

The midplane *interfacial surface area* is expected to directly correlate with the extent of interdigitation, as shown in Fig. 1. When this quantity is normalized by the bilayer area projected onto the *xy* plane, the resulting dimensionless quantity—which we term *nISA*— represents the fractional increase in the interfacial surface area due to interdigitation. We calculated the interfacial surface area between top and bottom leaflet lipids for each frame as follows. All pairs of neighboring atoms (defined as those whose Voronoi cells share a face) in which the two atoms also belonged to different leaflets were identified, and the areas of their shared faces summed to give the total interfacial surface area. This area was then divided by the cross-sectional area of the box in the frame to yield *nISA*, i.e.,

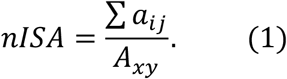

In Eq. 1, *a*_*ij*_ is the area of the face *f*_*ij*_ common to the Voronoi cells for atom *i* in the top leaflet and atom *j* in the bottom leaflet, and *A*_*xy*_ is the lateral area of the simulation box. The interleaflet contact surface can be visualized in 3D by plotting a mesh grid of the vertex coordinates of the common faces (Fig. 2A), or in 2D by projecting the coordinates onto the *xy* plane and using grayscale to represent the height in the *z*-dimension (Fig. 2B).

**Figure 1.**
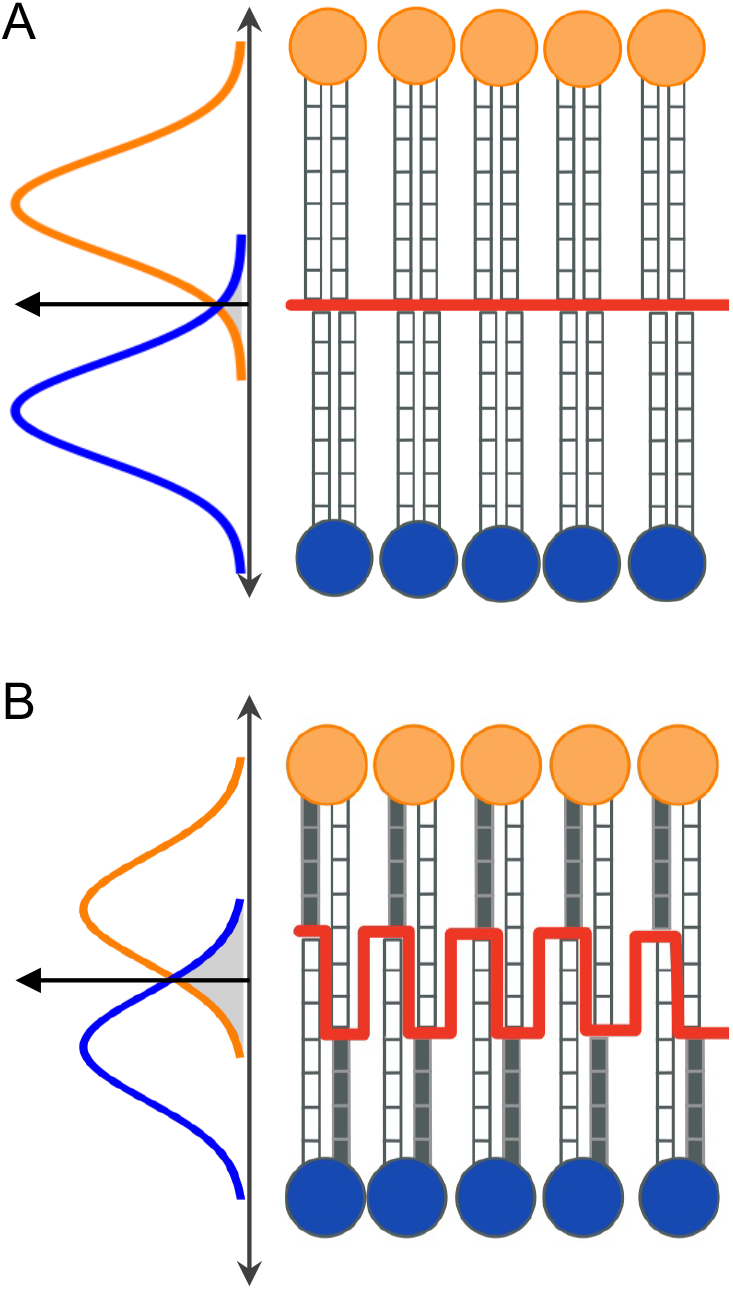
Schematic illustration of interdigitation quantified by the normalized interfacial surface area (nISA) and the Interdigitation tool in MEMBPLUGIN (memIT). (A) nISA is calculated as the midplane surface area (in red) divided by the lateral bilayer area. In the absence of interdigitation, the interfacial surface area per lipid is equal to the average area per lipid and nISA = 1. memIT, quantified by the overlap of the total mass density profiles of the top (orange) and bottom (blue) leaflets, is also shown as a gray shaded area. (B) Increased interdigitation in a mixed-chain bilayer results in a greater mass density overlap and a more dimpled midplane surface, leading to an increase in both memIT (larger gray area) and nISA (larger interfacial surface).

**Figure 2.**
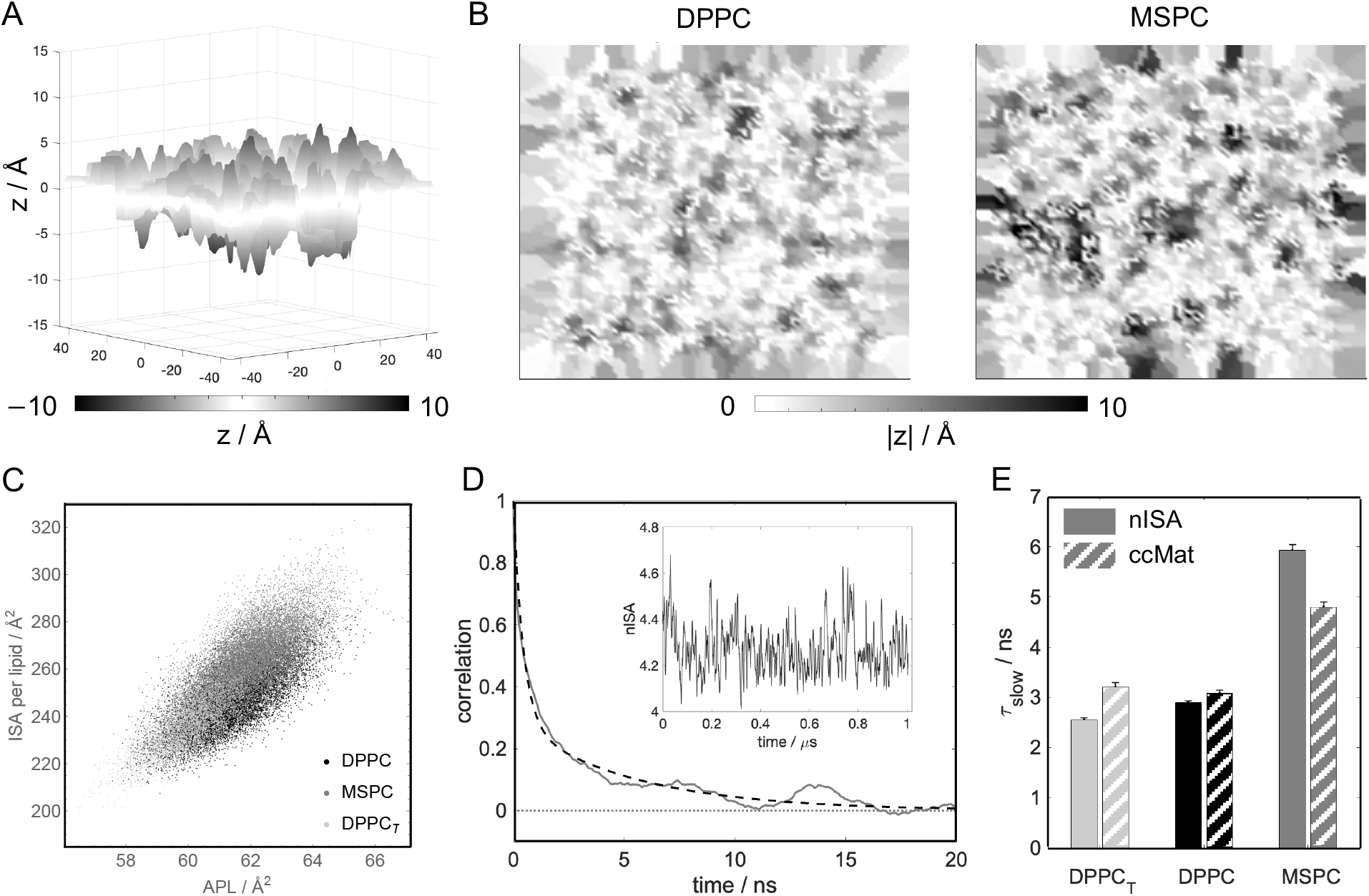
Interdigitation quantified with nISA. (A) 3D visualization of the interleaflet contact surface of DPPC from a representative trajectory frame. The surface is defined as the set of Voronoi cell faces shared by neighboring interleaflet atom pairs as described in Methods. (B) Contact surfaces shown as 2D intensity maps of |*z*| where darker shades correspond to greater interdigitation of acyl chains for DPPC and MSPC. (C) Interfacial surface area per lipid is correlated with the average area per lipid across all bilayers (correlation of 0.75). (D) Time autocorrelation function of nISA calculated for MSPC (solid line) and its best fit to a double exponential function (dashed line). Inset shows the raw data as a function of time. (E) Slow correlation times for nISA and ccMat for the three bilayers. Data and fits are shown in Table S2 and Fig. S4.

### Carbon Contact Matrix (ccMat)

In addition to the interfacial surface area, interdigitation in a bilayer can also be quantified by the number of direct interactions between acyl chain carbons of lipids in opposing leaflets. We counted these interactions by using a cutoff distance of 4 Å to define atomic contact.^41^ A matrix was constructed in which each row and column represent distinct chain carbons of the top and bottom leaflet lipids, respectively, and each matrix element is the time averaged number of contacts between these carbon atoms. The data is visualized as a heatmap in which the color intensity of the pixel represents the average number of opposing leaflet carbon pair interactions.

### Density Overlap Matrix (doMat)

Interactions between lipid carbon atoms from opposing leaflets can also be quantified from the overlap of the carbons’ number density profiles (NDPs). We used VMD’s Density Profile tool to calculate the NDP for each carbon of the top and bottom leaflet lipid chains. The overlap area, Ω_*ij*_, was then calculated as:

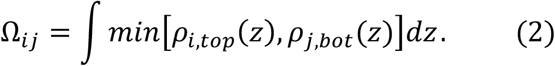

In Eq. 2, *ρ*_*i,top*_(*z*) is the NDP for carbon *i* in the top leaflet, *ρ*_*j,bot*_(*z*) is the NDP for carbon *j* in the bottom leaflet, and the integral is computed over all *z* (in practice, the lower and upper bounds of the simulation box in the *z*-dimension). A matrix similar to that calculated for the ccMat method was then constructed, with each entry corresponding to the overlap for one carbon pair. The sum of intensities in the four quadrants and the whole matrix provide alternative metrics for the extent of interdigitation.

Additional methodological details are described in Supplemental Information.

## RESULTS AND DISCUSSION

To investigate interdigitation, we simulated two bilayers made of lipids with either matching (DPPC) or mismatched (MSPC) chain lengths and shown recently in scattering experiments to have substantially different extents of interdigitation.^21^ As an additional control, we simulated DPPC under a constant negative surface tension (DPPC_T_), which compresses the bilayer area (Table S1) and should decrease interdigitation. We first used the standard method for quantifying interdigitation by calculating the overlap of the mass densities of all atoms in the two leaflets (memIT, see Supplemental Methods and Table 1). This calculation confirmed that MSPC chains have the highest degree of interdigitation, followed by DPPC and DPPC_T_, as expected (Fig. S1). The trend is consistent with results obtained from analysis of small-angle scattering data, which showed a 28% increase in the overlap of terminal methyl carbons for MSPC compared to DPPC.^21^

**Table 1.** Interdigitation quantified by various methods: memIT was calculated using all leaflet atoms; nISA is the fractional increase of the lateral bilayer area at the interfacial surface; ccMat corresponds to the average number of total trans-leaflet carbon-carbon contacts per frame; and doMat is the sum of the overlap number density integral of all trans-leaflet carbon-carbon pairs (see Methods and Supplemental Methods). Errors were calculated with block averaging as detailed in Supplemental Methods.

| System | memIT<br>(dimensionless) | nISA<br>(dimensionless) | ccMat<br>(contacts/frame) | doMat ($\text{\AA}^{-2}$) |
| --- | --- | --- | --- | --- |
| DPPC | $0.282 \pm 0.0013$ | $4.03 \pm 0.010$ | $270 \pm 1.1$ | $0.75 \pm 0.009$ |
| MSPC | $0.324 \pm 0.0014$ | $4.26 \pm 0.010$ | $342 \pm 1.5$ | $0.88 \pm 0.011$ |
| DPPC<sub>T</sub> | $0.270 \pm 0.0019$ | $3.94 \pm 0.011$ | $257 \pm 1.5$ | $0.69 \pm 0.024$ |

Next, we used 3D Voronoi tessellation to examine the interface that separates the two leaflets. A representative snapshot of the interfacial surface reveals that MSPC lipid chains penetrate deeper into the opposing leaflet compared to the chains of DPPC (Fig. 2A−B). We also calculated a dimensionless normalized interfacial surface area, nISA, from the 3D Voronoi analysis (Eq. 1, Table 1). In the DPPC bilayer, nISA was 4.03 ± 0.010, indicating that the interfacial surface area per lipid is four times larger than the average area per lipid projected onto the *xy* plane, consistent with the three-dimensional nature of the true interleaflet contact surface compared to idealized interactions at a flat bilayer midplane. In the MSPC bilayer, nISA was 4.26 ± 0.010, indicating a greater extent of interdigitation compared to DPPC in agreement with the trend seen in memIT values (Table 1, Fig. S1). Similarly, the compressed DPPC_T_ bilayer had the smallest nISA of 3.94 ± 0.011, as expected. The data also allows for extraction of the specific contributions of the sn-1 and sn-2 chains to the nISA surface, revealing their relative abundance in cross-leaflet interactions (Fig. S2). Furthermore, the high temporal resolution of the simulations enables an examination of interdigitation dynamics. For example, we find that the interfacial surface area is dynamically correlated with the projected box area (Fig. 2C), confirming that increased interdigitation leads to a decrease in thickness and increase in bilayer area (with area and thickness being strongly correlated across all bilayers, Fig. S3). Analysis of the nISA time autocorrelation function (Fig. 2D) reveals the presence of slow- and fast-decaying processes that are 2 to 4 times longer for MSPC compared to DPPC and DPPC_T_, indicating longer-lived interleaflet contacts for the more interdigitated MSPC membrane (Table S2, Figs. 2E, S4A−C).

Applying a conceptually different approach, we then compared the number of pairwise contacts between acyl chain carbon atoms of lipids in opposing leaflets by calculating a carbon-contact matrix, ccMat (Fig. 3A). The MSPC bilayer had on average ∼27% more cross-leaflet carbon-carbon contacts per frame than DPPC and ∼33% more contacts than DPPC_T_ (Table 1, Fig. S5A). For DPPC, interleaflet contacts were greatest for the terminal methyl carbons and decreased systematically with increasing distance from the terminal methyls, with only minor differences in sn-1/sn-1, sn-2/sn-2, and sn-1/sn-2 interactions. In contrast, ccMat for MSPC revealed that the largest number of interleaflet contacts occurs between the terminal methyl carbons of different chains (i.e., sn-1/sn-2), followed by the terminal methyl carbons of the longer chain (sn-2/sn-2). The asymmetry in the MSPC matrix is consistent with the longer chain extending deeper into the opposing leaflet and engaging in a greater total number of contacts with the longer chain of the lipids therein (Fig. 3A, bottom right quadrant), as also observed in the nISA analysis of specific chain interactions (Fig. S2). Comparison of the interdigitation dynamics quantified with ccMat across all bilayers shows the same trend as that of nISA (Table S2, Figs. 2E, S4 D-F).

**Figure 3.**
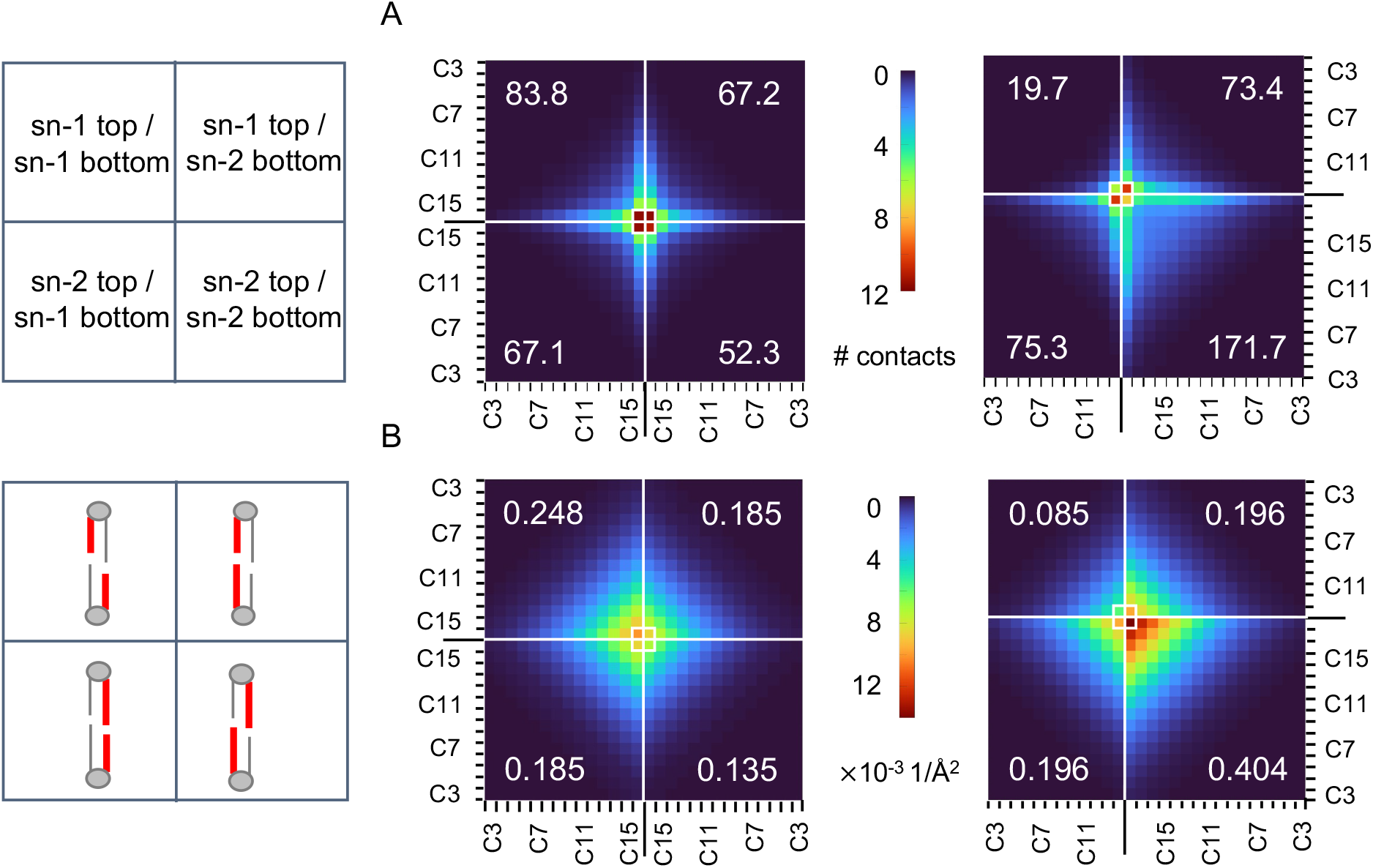
Matrix-form quantification of lipid interdigitation in DPPC (left) and MSPC (right) bilayers. Quadrants correspond to interactions between carbon atoms in the sn-1 and sn-2 chains of opposing leaflets as indicated in the cartoons on the left. (A) Carbon contact matrices (ccMat) where each matrix element represents the time-averaged number of close contacts for the specific interleaflet carbon pair. (B) Atomic density overlap matrices (doMat) where each matrix element represents the overlap of time-averaged atomic number density profiles for the interleaflet carbon pair calculated from Eq. 2. The mean quadrant sums are indicated in the corresponding four corners of each matrix and their errors are listed in Table S1.

The ccMat analysis uses an arbitrary cutoff distance for interatomic contacts. To circumvent this requirement, we also quantified interdigitation from the overlap of the number density profiles of individual cross-leaflet carbon pairs, an approach that we termed doMat. Fig. 3B shows doMat for MSPC and DPPC, again revealing a relatively symmetric matrix for DPPC and an asymmetric matrix for MSPC, consistent with the differences in chain mismatch for these lipids. Qualitatively, doMat features are similar to ccMat, with the overlap of cross-leaflet carbon atoms systematically decreasing with distance from the tail ends (Fig. 3B). While sn-1/sn-2 interactions are similar in the two bilayers, DPPC has a more than 3-fold larger overlap of sn-1/sn-1 number densities compared to MSPC (and more than 4-fold increase in close contacts from the ccMat analysis, Fig. 3A), indicating that cross-leaflet interactions between two shorter chains in the sn-1 positions are infrequent for chain-asymmetric lipids. Furthermore, consistent with ccMat, doMat for MSPC shows extensive overlap between carbons in the sn-2 chains of opposing leaflet lipids (Fig. 3B, lower right quadrant). Interestingly, the sn-2/sn-2 terminal carbon interactions are more pronounced than the sn-1/sn-2 interactions in doMat, in contrast to the trend in ccMat. This can be explained by the larger conformational space sampled by the longer stearoyl chains, which results in atomic number densities extending further into the opposing leaflet and a concomitant increase in interleaflet carbon overlap. The sum of the doMat matrix elements showed 17% and 27% more carbon-carbon overlap in the MSPC bilayer compared to DPPC and DPPC_T_, respectively (Table 1, Fig. S1, S5B).

The code for the three methods, as well as detailed instructions and examples, have been deposited on Zenodo (10.5281/zenodo.14508993) and should be straight-forward to implement. The provided implementation uses a structure .psf file and a simulation trajectory in .dcd format and is based on mostly freely available software packages (the only exception is the analysis in MATLAB which can be easily ported to e.g. python). Table S3 lists the required software and libraries. Of the three methods, nISA takes the longest to compute (∼1 min/frame for bilayers with 200 lipids), however the trajectory can be analyzed in multiple blocks in parallel for more efficient performance.

## CONCLUSION

The modes of interleaflet interaction in a lipid bilayer are tightly coupled to properties such as membrane thickness and local leaflet elasticity.^42, 43^ Here, we present three methods for quantitatively characterizing the physical contact between the leaflets as manifested by the degree of lipid interdigitation. While some of them, for example the number of cross-leaflet contacts and overlap of number densities, can be calculated with existing tools for specific atom selections (as in memIT), analysis of the trends along the entire lipid chains (as in ccMat and doMat) provides additional details and allows for a more comprehensive representation of lipid interdigitation. Furthermore, quantification of the entire spatially-resolved midplane surface and its time evolution enables in-depth analysis of the dynamics of all direct leaflet-leaflet interactions, something that existing methods do not provide.

While the three methods can in principle be applied to bilayers of varying sizes, it is important to consider their limitations. One advantage of ccMat is that it is based on local analysis of interleaflet carbon contacts and therefore should not be affected by membrane undulations that are often observed in larger bilayers. In contrast, doMat relies on number densities calculated in flat slabs along the bilayer normal making the results sensitive to membrane curvature. As currently implemented, nISA will also be affected by undulations since the quantity is normalized by the instantaneous (projected) bilayer area (Eq. 1). Normalizing the interfacial surface area by the number of lipids instead, can help alleviate this problem.

The computational results obtained from these methods can be validated by ensuring consistency with experimental measurements when available; for example, lipid volumes calculated from the 3D Voronoi tessellation protocol can be compared to those obtained from densitometry measurements.^44, 45^ Reliable simulation findings can be used to investigate the energetics of leaflet-leaflet interactions and their contribution to bilayer mechanics, to examine the relationship between interdigitation and other local lipid dynamics like lipid protrusion, and to generate testable hypotheses for the mechanisms of interleaflet coupling. The results can thus provide valuable insights for general structure-property relations in lipid bilayers.

## DATA AND SOFTWARE AVAILABILITY

The simulation trajectories and analysis scripts are available at 10.5281/zenodo.14508993.

## AUTHOR INFORMATION

### Authors

**Emily Chaisson** – Department of Chemistry, University of Tennessee Knoxville, Knoxville, TN 37996, USA

### Notes

The authors declare no competing financial interest.

## AUTHOR CONTRIBUTIONS

E.H.C. performed simulations and analysis. F.A.H. performed analysis. F.A.H. and M.D. designed the research. E.H.C., F.A.H. and M.D. wrote and edited the manuscript.

## ACKNOWLEDGMENTS

This research was supported by NIH grant R01 GM138887 (to FAH) and the SciLifeLab & Wallenberg Data Driven Life Science Program grant: KAW 2024.0159 (to MD).

## TOC Graphic

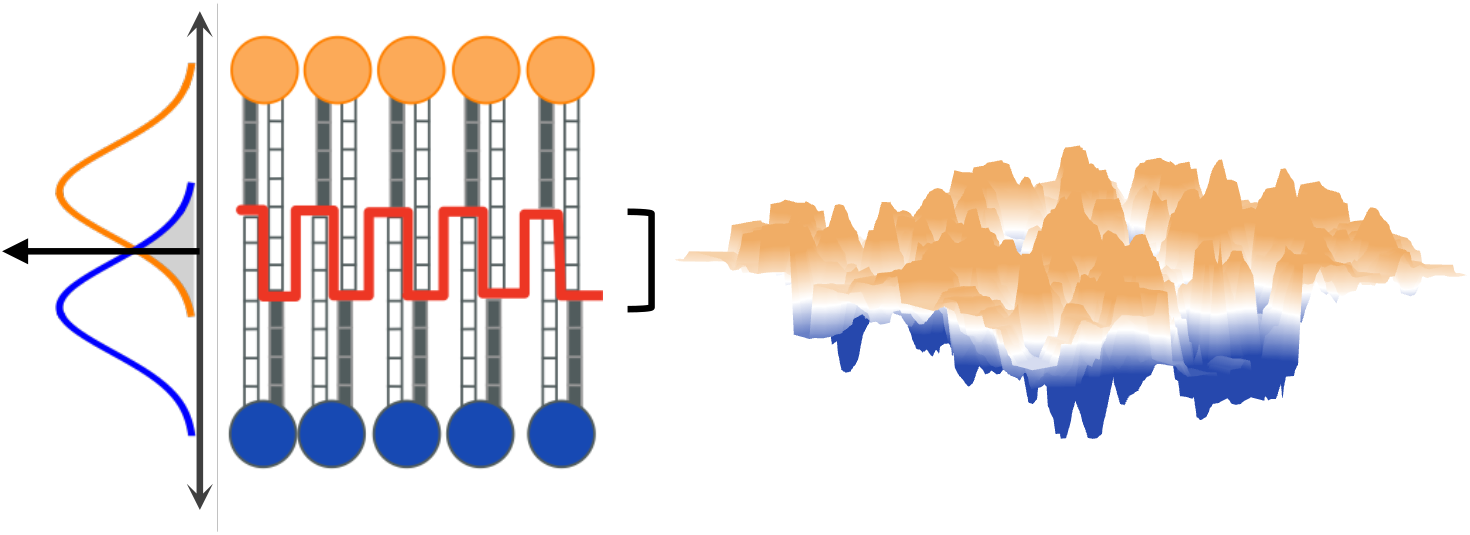

