## Supplemental Material for "QUANTIFYING ACYL CHAIN INTERDIGITATION IN SIMULATED BILAYERS VIA DIRECT TRANSBILAYER INTERACTIONS"

The document includes supplemental methods, 3 tables and 5 figures.

SUPPLEMENTAL METHODS

*MEMBPLUGIN Interdigitation Tool (memIT).* The MEMBPLUGIN[1] package for VMD has a *Lipid Interdigitation* tool that provides three measures of interdigitation: $I_p$, $w_p$, and $I_C$. [1] $I_p$ and $w_p$ are the fraction and width (respectively) of the mass density overlap calculated for a user-specified selection of atoms from the two bilayer leaflets. $I_C$ is the fraction of inter-leaflet atomic contacts calculated from all lipid atoms with a cutoff distance of 4 Å. For comparison with the new methods, we used $I_p$ and performed the analysis on all atoms in the leaflets, including hydrogens.

*Error analysis*. For all measured quantities, reported uncertainties are standard errors of the mean calculated by block averaging to properly account for time correlations; an excellent description of this procedure can be found in a recent methods paper from Foley and Deserno[2]. Briefly, consecutive measurements in the time series were iteratively blocked into groups of two and averaged, with the standard error calculated at each blocking iteration. A plot of the standard error vs. blocking iteration shows an initial increase and eventual plateau once the block size is large enough for successive values to be uncorrelated; the uncertainties we report are the standard error at the plateau region. Since the doMat analysis already requires frame averaging, uncertainty was estimated as the standard error of doMat values calculated separately for consecutive blocks of ~1000 frames (a total of 5-10 blocks depending on the trajectory length). For comparing differences in the means of two distributions, p-values were calculated from the Student's *t* distribution with an effective number of degrees of freedom implied by the block averaging procedure.

SUPPLEMENTAL TABLES

**Table S1.** Interdigitation and structural parameters of simulated bilayers. Listed are the results from memIT, nISA, ccMat and doMat (both the total matrix sums and individual quadrant sums), the average area per lipid and average phosphate-to-phosphate distance (thickness). Errors are calculated from block averaging as described in Supplemental Methods.

| | $DPPC_T$ | DPPC | MSPC |
|---|---|---|---|
| memIT / dimensionless | $0.270 \pm 0.0019$ | $0.282 \pm 0.0013$ | $0.324 \pm 0.0014$ |
| nISA / dimensionless | $3.94 \pm 0.011$ | $4.03 \pm 0.010$ | $4.26 \pm 0.010$ |
| ccMat / contacts per frame | $257 \pm 1.5$ | $270 \pm 1.1$ | $342 \pm 1.5$ |
| ccMat quadrant 1 | $64.1 \pm 0.4$ | $67.2 \pm 0.4$ | $73.4 \pm 0.6$ |
| ccMat quadrant 2 | $80.6 \pm 0.7$ | $83.8 \pm 0.4$ | $19.7 \pm 0.6$ |
| ccMat quadrant 3 | $63 \pm 1$ | $67.1 \pm 0.3$ | $75.3 \pm 0.7$ |
| ccMat quadrant 4 | $48.7 \pm 0.6$ | $52.3 \pm 0.4$ | $171.7 \pm 0.7$ |
| doMat / $Å^{-2}$ | $0.69 \pm 0.024$ | $0.75 \pm 0.009$ | $0.88 \pm 0.011$ |
| doMat quadrant 1 | $0.165 \pm 0.007$ | $0.185 \pm 0.002$ | $0.196 \pm 0.003$ |
| doMat quadrant 2 | $0.230 \pm 0.007$ | $0.248 \pm 0.003$ | $0.085 \pm 0.002$ |
| doMat quadrant 3 | $0.172 \pm 0.006$ | $0.185 \pm 0.002$ | $0.196 \pm 0.003$ |
| doMat quadrant 4 | $0.124 \pm 0.005$ | $0.135 \pm 0.002$ | $0.404 \pm 0.004$ |
| APL / $Å^2$ | $60.1 \pm 0.11$ | $61.42 \pm 0.071$ | $62.04 \pm 0.073$ |
| PP-dist / Å | $40.14 \pm 0.058$ | $39.50 \pm 0.036$ | $39.20 \pm 0.039$ |

**Table S2.** Correlation times of interdigitation dynamics. Fluctuations in nISA and ccMat were fit with a double exponential decay (Fig. S4). Shown are the respective correlation times and errors calculated from the fits (using Mathematica's NonlinearLeastSquares function).

| bilayer | nISA / ns | | ccMat / ns | |
|---|---|---|---|---|
| | $\tau_{\mathrm{fast}}$ | $\tau_{\mathrm{slow}}$ | $\tau_{\mathrm{fast}}$ | $\tau_{\mathrm{slow}}$ |
| MSPC | $0.481 \pm 0.008$ | $5.93 \pm 0.11$ | $0.082 \pm 0.003$ | $4.8 \pm 0.1$ |
| DPPC | $0.141 \pm 0.007$ | $2.90 \pm 0.03$ | $0.036 \pm 0.003$ | $3.08 \pm 0.065$ |
| $DPPC_T$ | $0.121 \pm 0.007$ | $2.55 \pm 0.04$ | $0.039 \pm 0.003$ | $3.2 \pm 0.1$ |

**Table S3.** Software and libraries used in the implementation of the three interdigitation methods.

| Method | Software | Special libraries | Source/download |
|---|---|---|---|
| all | VMD | | https://www.ks.uiuc.edu/Development/Download/download.cgi?PackageName=VMD |
| | MATLAB | | https://mathworks.com/help/install/install-products.html |
| nISA | voro++ | | http://math.lbl.gov/voro++/ |
| | python3 | pandas<br>csv | https://pypi.org/project/pandas/<br>https://docs.python.org/3/library/csv.html |
| ccMat | VMD | tcllib | https://wiki.tcl-lang.org/page/Tcllib+Installation |
| doMat | VMD | Density Plugin | https://github.com/giorginolab/vmd_density_profile |

SUPPLEMENTAL FIGURES

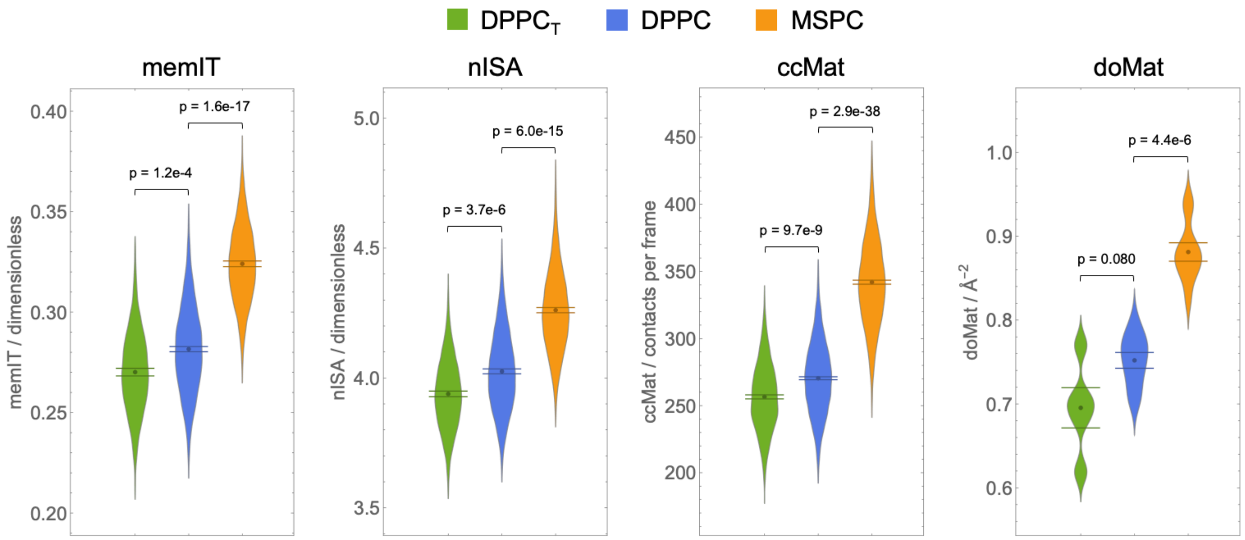


**Figure S1.** Interdigitation in simulated bilayers quantified with different methods. Shown for each method are: the spread in the values from individual frames (for memIT, nISA and ccMat) or consecutive blocks of ~1000 frames (for doMat); their mean and effective standard error of the mean calculated from block averaging as detailed in Supplemental Methods; and the *p*-values for comparisons of the results for $DPPC_T$ and MSPC against those for the DPPC bilayer.

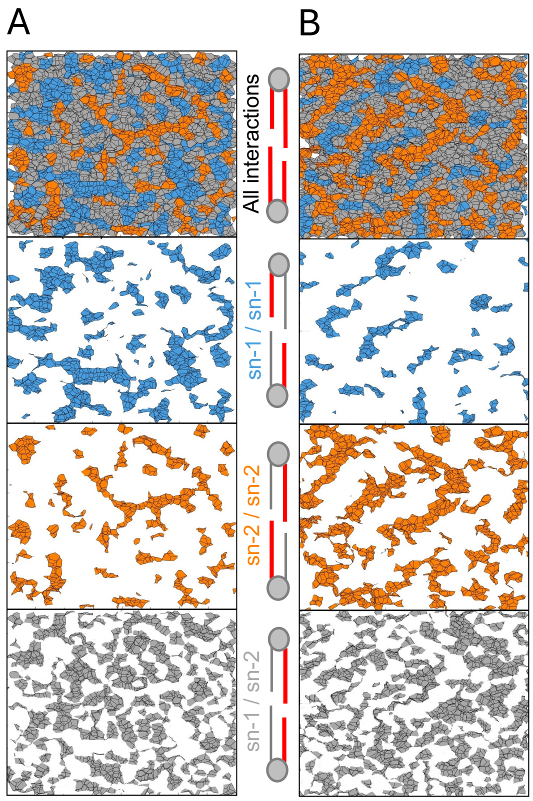


**Figure S2.** Maps of shared Voronoi faces between atoms in opposing leaflets in DPPC (A) and MSPC (B) from the same trajectory frames as in Fig. 2. Different maps correspond to (from top to bottom): all atom pairs, sn-1/sn-1 pairs, sn-2/sn-2 pairs, and sn-1/sn-2 pairs. Each map is $80 \times 80$ Å.

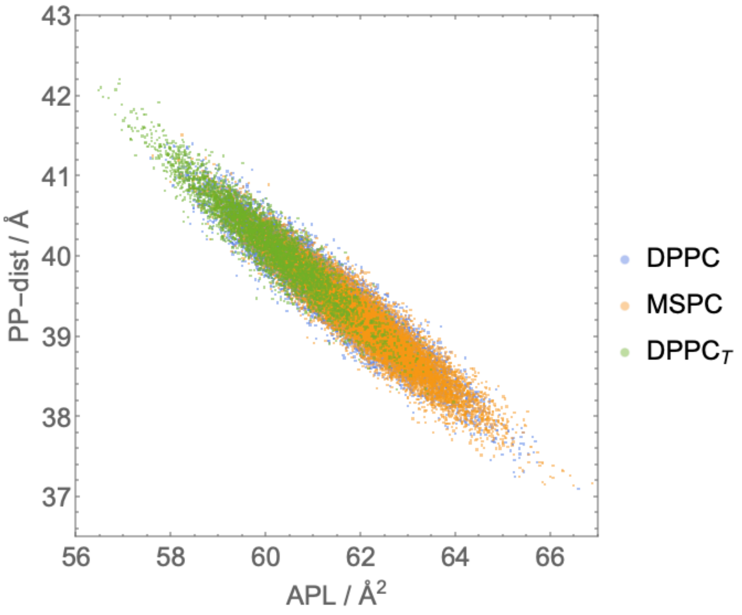


**Figure S3**. Correlation between bilayer thickness (phosphate-to-phosphate distance, PP-dist) and area per lipid (APL) in the three simulated bilayers. Each data point represents a single frame from the corresponding trajectory.

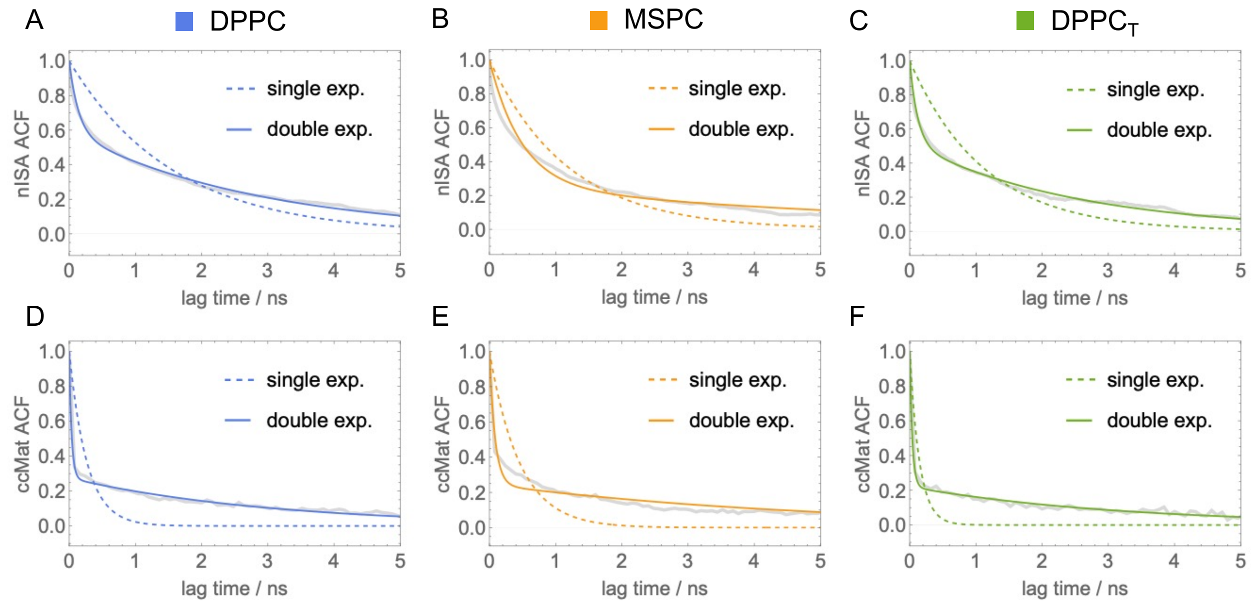


**Figure S4.** Autocorrelation functions of nISA (top) and ccMat (bottom) for each simulated bilayer (gray). Shown on each plot are the corresponding best fits with a single exponential (dashed color line) vs double exponential (solid color line) decay. The resulting correlation times from the double exponential fits are listed in Table S2.

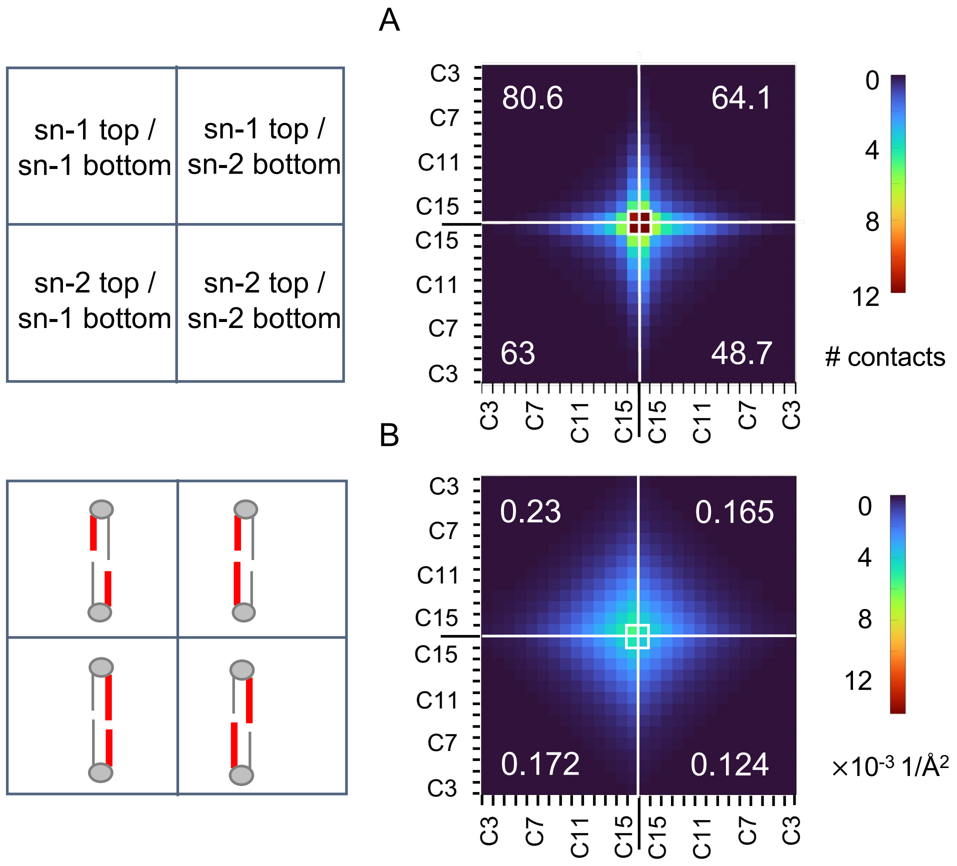


**Figure S5.** Interdigitation in the DPPC$_T$ bilayer quantified with ccMat (A) and doMat (B). Shown are the average results from each method as matrices following the format of Fig. 3 in the main text. Errors for the quadrant sums are listed in Table S1.
